# Physical constraints during Snowball Earth drive the evolution of multicellularity

**DOI:** 10.1101/2023.12.07.570654

**Authors:** William W. Crockett, Jack O. Shaw, Carl Simpson, Christopher P. Kempes

## Abstract

Molecular and fossil evidence suggest that complex eukaryotic multicellularity evolved during the late Neoproterozoic era, coincident with Snowball Earth glaciations, where ice sheets covered most of the globe. During this period, environmental conditions—such as sea water temperature and the availability of photosynthetically active light in the oceans—likely changed dramatically. Such changes would have had significant effects on both resource availability and optimal phenotypes. Here, we construct and apply mechanistic models to explore (i) how environmental changes during Snowball Earth and biophysical constraints generated selective pressures and (ii) how these pressures may have had differential effects on organisms with different forms of biological organization. By testing a series of alternate—and commonly debated—hypotheses, we demonstrate how multicellularity was likely acquired differently in eukaryotes and prokaryotes due to selective differences in the biophysical and metabolic regimes they experience: decreasing temperatures and resource-availability instigated by the onset of glaciations generated selective pressures towards smaller sizes in organisms in a diffusive regime and towards larger sizes in motile heterotrophs. These results suggest that changing environmental conditions during Snowball Earth glaciations gave multicellular eukaryotes an evolutionary advantage, paving the way for the complex multicellular lineages that followed.

## I. Introduction

A fundamental focus of biology is understanding the vast range of body sizes and the associated diversity in the number of levels of hierarchical organization [1, 2]. Each new level of organization is typically associated with a major event in evolutionary history that changed the state of the evolutionary game. By adding a new hierarchical level to the organization of organisms, these major transitions in individuality added new niches to the ecosystem (e.g., trophic) and introduced new phenotypes. Such transitions include the origin of cells, eukaryotes, multicellularity, and colonial and social organisms. The insight that these transitions share evolutionary processes involved in the emergence of a new level of organization has proven to be a powerful research program (see [1, 3–5] for comprehensive reviews of the topic).

However, it is challenging to understand certain transitions, such as multicellularity, because of the large number of independent origins, the fact that eukaryotes and prokaryotes both evolve multicellular forms, and the lack of substantial fossil and molecular evidence [6, 7]. The evolution of multicellularity stands as one of the most pivotal milestones in the history of life on Earth as it revolutionized biological organization and paved the way for the diversity of macro-scale organisms we observe today. Its emergence allowed for specialized cells to cooperate, leading to the development of complex tissues, organs, and organ systems. This enhanced complexity further facilitated the evolution of complex organisms with more sophisticated behaviors enabling adaptation to a wide range of environments and the exploitation of new ecological niches and new biological scales. Multicellularity laid the foundation for the diverse and interconnected web of life that shapes our planet’s ecosystems today.

Fossil and molecular evidence indicate that complex multicellularity originated and proliferated during the Neoproterozoic era (1,000 to 541 Ma) [8, 9]. Previous work commonly proposed that this evolution was connected to an increase in oxygen levels that removed a physical constraint on size. However, recent work suggests that sponges, a likely morphology for the last common metazoan ancestor, can survive oxygen levels as low as those present during the Neoproterozoic era [10], suggesting that low oxygen levels may not have been a physical constraint preventing the emergence of multicellular eukaryotes. Furthermore, other work suggests that the evolution of more complex eukaryotes including multicellular organisms could have led to ocean oxygenation [11] (as opposed to the other way around), and we know that multicellular eukaryotes can cope with low oxygen given that it is likely that the sea floor was anoxic when the first undisputed metazoan fossils appear in deep water [12–14]. If the appearance of multicellularity was not caused by changing oxygen levels, an alternative mechanism for why multicellular eukaryotes emerged during this period is needed.

Extreme glaciations during the Cryogenian period (∼ 720 − 635 Ma), a phenomenon commonly referred to as Snowball Earth, led to a radical transformation of the Earth’s climate and oceans [15]. Across two major glaciations, lasting almost 50 million years, glaciers appear to have reached the equator, although there is still debate over the extent of coverage [16, 17]. The global glaciations resulted in the widespread freezing of the planet’s surface, severely restricting the availability of light and nutrients to depths below. Prior to Snowball Earth, simulations suggest the ocean was relatively warm, with surface water temperatures reaching 30 °C at the equator [18]. However, depending on the severity of glaciations, temperatures likely dropped to between -4 °C and 4 °C [17, 19]. Given that such extreme conditions persisted for many tens of millions of years, it is important to understand how these conditions would have affected the ability of single-celled organisms to survive and reproduce. Notably, fossil evidence does not indicate any significant extinctions [20, 21]. One potential means of success in these conditions may have been found in the formation of cooperative groups of cells in some lineages, which then could have lead to the emergence of multicellular life.

Recent work [22] suggests that the long-term loss of low-viscosity environments, instigated by decreasing ocean temperatures during the Cryogenian, generated selective pressures towards multicellularity in eukaryotes. This work suggests that adaptation to environmental conditions led to larger sizes and speeds only accessible through multicellularity to exploit limited resources and satisfy metabolic needs during Snowball Earth’s high-viscosity regimes. Following the cessation of glaciation and the return of low-viscosity environments these newly evolved multicellular taxa remained and proliferated.

Beyond the viscosity shifts associated with the much lower temperatures of Snowball Earth there are many other physical, physiological, and ecological changes expected during this interval (e.g., [17, 23–25]). For example, the accumulation of significant sea ice likely decreased light flux to the ocean and decreased the terrestrial nutrient run-off [16, 17]. Ecological and biogeochemical features associated with sinking, remineralization, predation, and the size distribution of organisms are all also expected to shift in this new environment.

For an organism to survive it must be able to access enough nutrients to satisfy metabolic demands. Several factors can be altered and integrated to allow an organism to increase nutrient capture, including metabolic rate, motility, and size. Given the existence of numerous optima, the specific combination of changes to metabolic rate, motility, and size is less important than the first-order need to acquire nutrients.

Because of the multiple contemporaneous origins of eukaryotic multicellularity an environmental driver is likely. However, an environmental driver can’t be universal because only a few of the many co-occurring eukaryotic lineages evolved multicellularity, such that the driver must also sort between adaptive strategies. An answer may be found if there are competing biophysical aspects that share a common cause. Cold conditions during Snowball Earth may provide such a cause, with effects on viscosity, diffusivity, and metabolic rates that lead to complex tradeoffs.

This paper presents analyses of mechanistic models for exploring interactions between the environmental changes associated with Snowball Earth, physical constraints on biological processes, and differential selective pressures between single-celled and simple multicellular organisms. First, we describe a global productivity model that suggests Snowball Earth’s changes in temperature and light availability generated a significant decrease in primary production. Second, based on this insight, we compare two models that describe how organisms with different biological organizations - a non-motile unicellular organism relying on diffusion, Fig. 1a, and a simple motile multicellular organism - are affected by the environmental changes predicted during Snowball Earth.

**Fig. 1.**
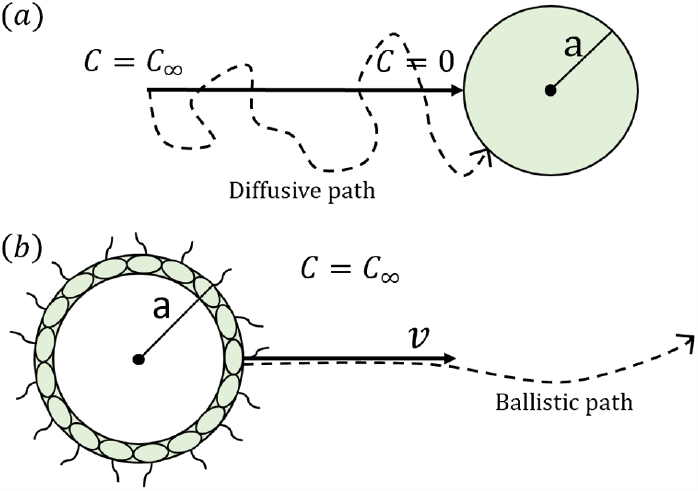
(*a*) Diagram of the non-motile diffusive cell. The spherical cell takes in all nutrients at the cell’s surface (*C* = 0), causing chemical resources (e.g. glucose) to diffuse toward the cell from far away (*C* = *C*_*∞*_). (*b*) Diagram of the motile choanoblastula. The organism is hollow with an outer radius *a*, and swims at a velocity *v*. The organism’s motility means it travels ballistically relative to its prey. Resource concentration is assumed to be constant (*C* = *C*_*∞*_).

For our multicellular organism we model a hypothetical and idealized ”choanoblastula” (Fig. 1b). The choanoblastula is heterotrophic, motile, and composed of a hollow-sphere of cells, such that it has similar morphology and physiology to the green algae genus *Volvox*, except that it does not photosynthesize. Something akin to this model organism may have existed during the Cryogenian, but would have been displaced by descendant lineages (e.g., metazoa).

Our results suggest differential responses to selective pressures: (i) for organisms operating in the diffusive regime, decreasing temperature and resource availability leads to a decrease in organismal size; and (ii) for motile heterotrophs with a simple multicellular morphology, environmental changes accompanying Snowball Earth selected for larger organisms.

## II. Methods

### A. Global Productivity Model

To understand the impacts that Snowball Earth had on eukaryotes and early metazoa, it is crucial to understand how the environmental changes impacted the broader ecosystem. A simple method to estimate the magnitude of these changes is to calculate the net primary productivity (NPP) as a function of temperature and intensity of photosynthetically active radiation (PAR) [26]:

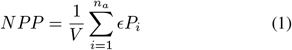

where *V* is the volume of water, *n*_*a*_ is the number of autotrophic cells, *ϵ* is the efficiency of production of organic matter, and *P*_*i*_ is the productivity of each autotrophic cell. The productivity of each autotroph can be modeled as a function of it’s metabolic rate and PAR. The metabolic rate is modeled using the metabolic theory of ecology (MTE) [27], which relates metabolism (*B*) to temperature (*T*) and organism mass (*M*_*i*_):

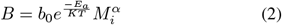

where *E*_*a*_ is the average activation energy of metabolic reactions, *b*_0_ is a constant, *K* is Boltzmann’s constant, and *α* is a power-law scaling term. The scaling term *α* is normally assigned a value of 3*/*4 for multicellular organisms, and 1 for single-celled eukaryotes [28, 29].

Productivity’s dependence on light intensity (*I*) is given by a Monod equation [30], where *K*_*I*_ is the half-saturating term. Combining the dependence of productivity on metabolic rate and light intensity results in the following expression [26]:

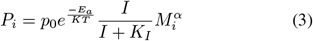

where *p*_0_ is a constant.

To model *n*_*a*_, the steady-state biomass model in [31] is employed. Assuming constant cell size, this model calculates the supported biomass under given nutrient flux conditions, allowing us to solve for the population carrying capacity for a given set of environmental conditions.

### B. Uptake-Metabolism Energy Balance

An energy balance was used to model the impact of changing temperature and resource concentration on organisms, where the rate of energetic resource uptake (*U*) must be greater than or equal to the rate of energy use in the organism’s metabolism (*B*):

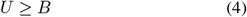

To understand how environmental changes altered optimal phenotypes, resource uptake and metabolism can be modeled as functions of temperature, resource concentration, and organismal traits (which are assumed to be generated from body size). Both rates depend on specific resource acquisition strategies and organism morphologies, two of which we explore here.

#### 1) The Non-motile Diffusive Cell

The modeled organism was inspired by smaller prokaryotes, with the following traits: single celled, non-motile, and reliant upon diffusion for uptake (Fig. 1a). Assuming that the cell takes up all resources at its surface, and that resource concentration approaches a constant (*C*_*∞*_) far away from the cell, we can solve the diffusion equation to obtain an equation for resource concentration:

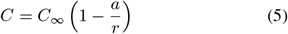

where *a* is the radius of the cell, and *C* is the nutrient concentration at some distance *r* from the cell’s center. The cell’s total resource influx can be determined by applying Fick’s Law of Diffusion [32] to calculate flux density and integrating it across the cell’s surface [33]:

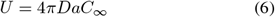

Here *D* is the diffusivity of the resource, which can be defined by the Stokes-Einstein equation [34]. Viscosity (*η*), can be modeled as a function of temperature using the Vogel–Fulcher–Tammann (VFT) equation [35]. Diffusivity is inversely proportional to this viscosity. By incorporating these physical models into the uptake model (Eq. 6) resource uptake for the diffusive cell is modeled as a function of temperature, resource concentration, and cell size:

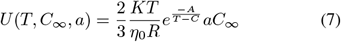

Equation 2 is used to model the metabolic rate of the diffusive cell [27]. Also, the conversion between volume and mass is approximated using a constant cell density. Using these definitions for resource uptake and metabolic rate in equation 4 and solving the inequality for organism radius (*a*) results in the model for the maximum diffusive cell size as a function of temperature and resource concentration:

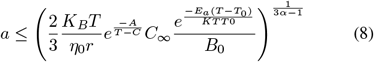

#### 2) The Motile Choanoblastula

The choanoblastula employs a different uptake strategy, and its morphology leads to a different metabolic scaling. The resource uptake rate is based on ballistic velocity of the organism, and its metabolism is based on the metabolic theory of ecology and an additional motility cost.

Due to the relative difference in velocity that arises from the choanoblastula’s motility, its uptake is ballistic rather than diffusive (Fig. 1) [36, 37]. In this case, the choanoblastula is colliding with its resource, causing resource uptake to scale with its cross-sectional area [38]:

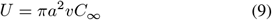

where *v* is the velocity of the choanoblastula relative to the resource. The velocity scales with organism radius and the viscosity of the surrounding fluid [39]. This is summarized in the generalized model [22]:

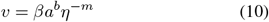

where *β* is a constant, and *b* and *m* are scaling coefficients. Estimates of *b* range from 0.5 to 1[22, 36, 40], and estimates of *m* range from 0.4 to 4 depending on the species [22], with a value of 1 found for *Chlamydomonas* [41]. Using the VFT equation to define viscosity and equation (10) to define velocity in equation (9) results in a model for ballistically motile resource uptake as a function of temperature and organism radius.

Organismal metabolism was modeled by employing the MTE (Eq. 2) to model basal metabolism with a motility cost. The basal metabolism scales with organismal mass, which is proportional to the number of cells in the organism. Due to its hollow-sphere morphology, the basal metabolic rate is proportional to organismal surface area:

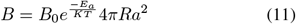

Assuming the organism exists at a Reynolds number less than 1 (i.e., where viscous forces of the fluid are dominant over inertial forces), the power it takes to maintain a velocity *v* through the fluid is given by Stokes’ Law [42], which, along with a coefficient of efficiency (*ϵ*), acts as the motility cost.

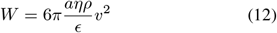

Incorporating each component of the model, the full energy balance becomes:

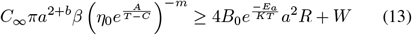

where *W*, the metabolic cost of motility, can be expanded using equations 10, 12 and the VFT equation to be a function of temperature and organism radius.

## III. Results

### A. Global Productivity Model

Four models of NPP were developed and analyzed under varying ecological and physiological responses to environmental changes (Fig. 2). Models were evaluated over the same range of temperature and PAR availability, but population size and producer size were either held constant or allowed to vary according to models.

**Fig. 2.**
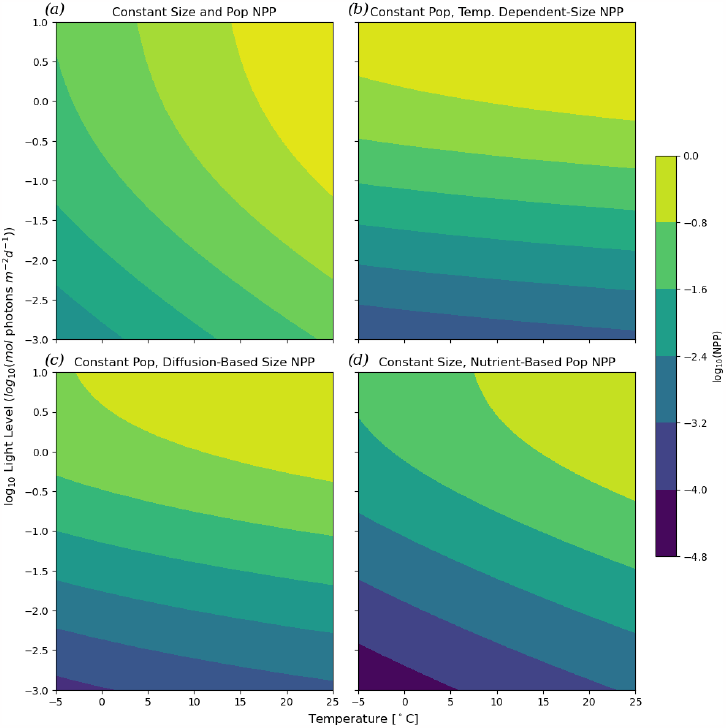
Contour-plots showing the log base 10 of net primary productivity (NPP) as a function of temperature [*°*C] (x-axis) and the relative log base 10 of photosynthetically-active light availability (y-axis). (*a*) Global NPP given a constant number of primary producers with constant mass. (*b*) Global NPP given constant number of primary producers, but their mass changes as a function of temperature based on the diffusion model (Eq. 8). (*c*) Global NPP given constant population size where the size of primary producers scales with the diffusion model (nutrient concentration is assumed to decrease with temperature, and is used to calculate producer size).(*d*) Global NPP where size is held constant, but population changes with temperature and limiting-nutrient concentration based on the Steady-State Biomass model in [31].

Under the best case, where primary producer mass and population size each remain constant with decreasing temperature and light, reduced metabolic rates lead to a 2 order-of-magnitude decrease in NPP (Fig. 2a).

In reality, most primary producers rely on diffusion to obtain the inorganic nutrients needed for growth. The diffusion model (Eq. 8) can be employed to consider how the primary producer’s size would have changed as temperature decreased. Assuming that both the concentration of inorganic nutrients and the number of primary producers are constant, introducing the temperature size dependence of the primary producers indicates that NPP would decrease by 2.5-3 orders-of-magnitude (Fig. 2b).

During the Cryogenian, environments capable of supporting life became more oligotrophic, reducing resource availability, and became eutrophic after melting [17, 43]. The impact of nutrient availability was incorporated into the NPP model by assuming that nutrient availability linearly decreases by half over the temperature interval. Nutrient availability could impact the size of primary producers (Fig. 2C) or the number of primary producers (Fig. 2D). Both cases lead to significant decreases in NPP, with an approximately 3.5 order-of-magnitude decrease for nutrient-limited cell size, and a 4.5 order-of-magnitude decrease for nutrient-limited population size.

Even when assuming resilient physiologies and ecosystems, decreased organic resource availability would have been a major environmental change for existing heterotrophic organisms.

### B. The Diffusive Cell

The non-motile diffusive cell’s (Eq. 8) dependence on temperature is two-fold: (i) the metabolic rate’s dependence on temperature and (ii) the uptake rate’s dependence on diffusivity and viscosity. The decrease in temperature that accompanied Snowball Earth caused an increase in viscosity accompanied by a decrease in diffusivity and nutrient uptake, but also led to a slower metabolic rate. Although uptake drops to less than half of its pre-Snowball Earth value, under an activation energy of 0.62 eV metabolic rate drops by nearly a factor of 10 (Fig. 6). The slow down in metabolic rate means that although the cell’s uptake slows, it is able to grow in size as temperature decreases.

Based on the results from the NPP calculation, it is important to consider a decrease in organic resource concentration in addition to temperature decrease during Snowball Earth. For the non-motile organism relying on diffusion, it must shrink in size, reducing its radius *a*, to adapt to lower resource availability (Fig. 3). Under the best supported parameter values, the model predicts a cell radius of approximately 10 *µ*m prior to Snowball Earth and a radius of approximately 300 nm during Snowball Earth. Importantly, we show that cell size changes are greatly impacted by the assumed value of average metabolic activation energy *E*_*a*_. This value influences how metabolism scales with temperature, impacting the relative change between uptake and metabolic rate (Fig. 6b). For all values of *E*_*a*_, there is a decrease in cell size as resource availability drops, but varying values of *E*_*a*_ can change the temperature dependence of diffusive cell size (Fig. 7). While the average metabolic activation energy determines the response to temperature, all diffusive organisms, regardless of *E*_*a*_, must have decreased in size to survive the Cryogenian period due to the decrease in resource availability.

**Fig. 3.**
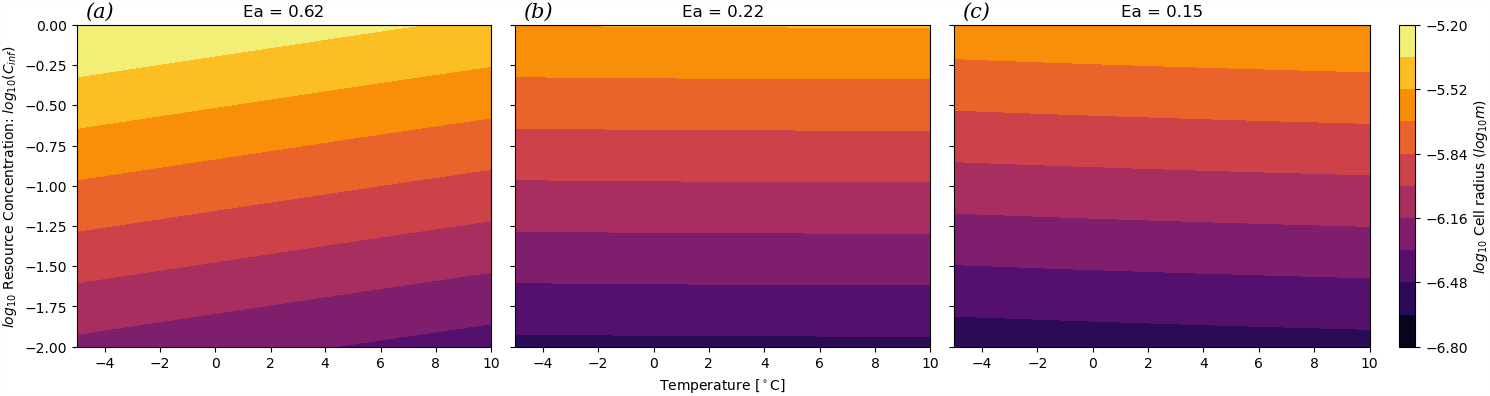
Contour-plot of the log of radius [log_10_(m)] of the diffusive cell as a function of temperature [*°*C] (x-axis) and relative resource concentration (y-axis). Each subplot shows the results under a different activation energy (*E*_*a*_).

**Fig. 4.**
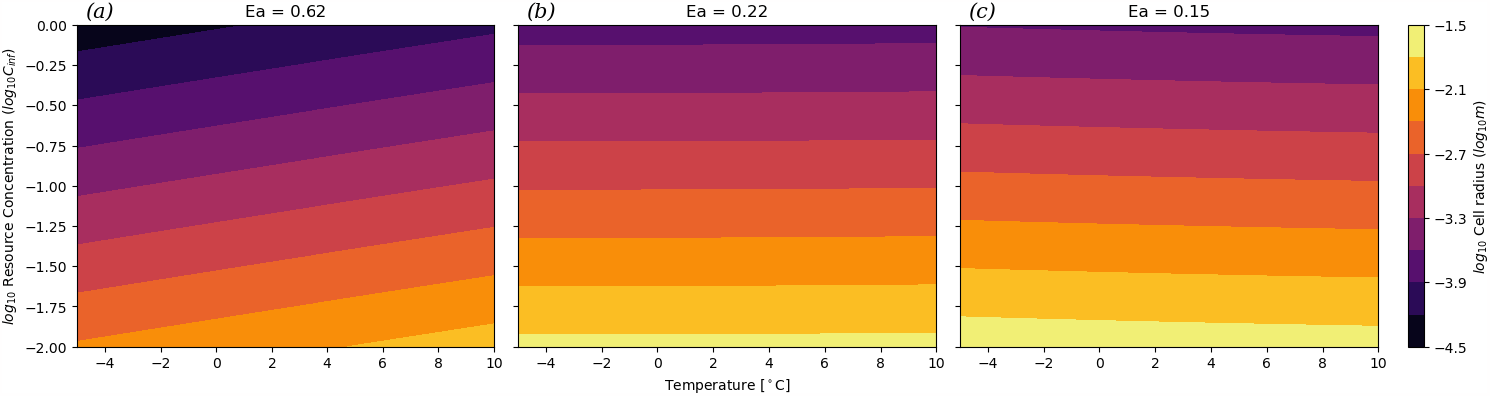
Contour-plot of the log of radius [log_10_(m)] of the choanoblastula as a function of temperature [*°*C] (x-axis) and relative resource concentration (y-axis). Each subplot shows the results under a different activation energy (*E*_*a*_). Plots are for *b* = 1.

### C. The Motile Choanoblastula

The choanoblastula’s motility introduces an additional temperature dependence to the energy balance due to the cost of motility’s dependence on viscosity (*η*) of water. However, the motility cost is relatively small compared to the basal metabolic cost and uptake rate, and therefore has a negligible effect (Fig. 5). Resource uptake scales with organism radius (*a*^1+2*b*^, where 0.5 *≤ b ≤* 1) more quickly than the metabolic rate, which scales with *a*^2^ due to cells only existing on the sphere’s surface. Because resource uptake scales at a higher rate, there exists a critical size where for smaller radii the metabolic rate is greater than the uptake rate, and for larger radii the uptake rate is greater than the metabolic rate (Fig. 5). This critical radius defines the minimum size of the organism for the given temperature and resource concentration, and is the solution to the energy balance in equation (13).

**Fig. 5.**
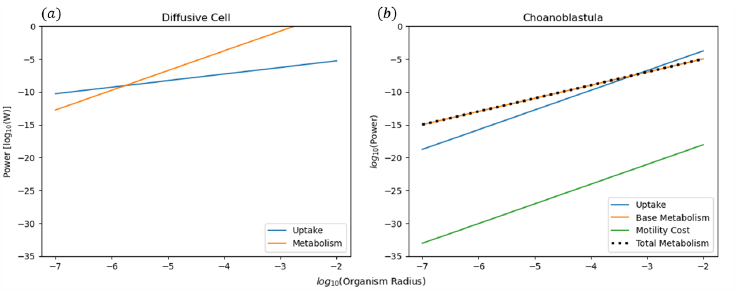
Energetic costs and nutrient uptake as a function of organism radius for the (*a*) diffusive cell and (*b*) the choanoblastula models (based on a temperature of 0 °C and nutrient concentration *C*_*∞*_ = 0.1).

**Fig. 6.**
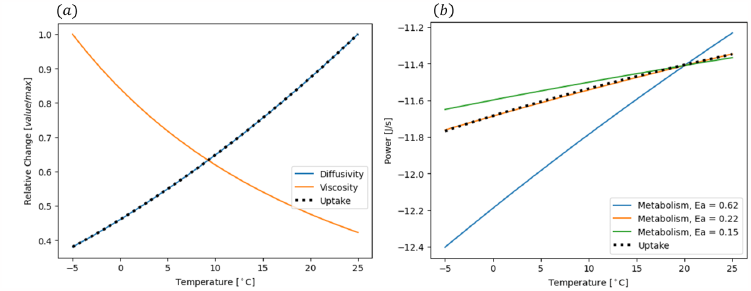
(*a*) Relative changes [*value/max*] in viscosity, diffusivity, and uptake rate for the diffusive cell as functions of temperature [^*°*^*C*]. (*b*) Value of uptake rate and metabolic rate [*log*_10_(*W*)] of the diffusive cell as functions of temperature [^*°*^*C*]. Metabolic rate is plotted for 3 different *E*_*a*_ values.

**Fig. 7.**
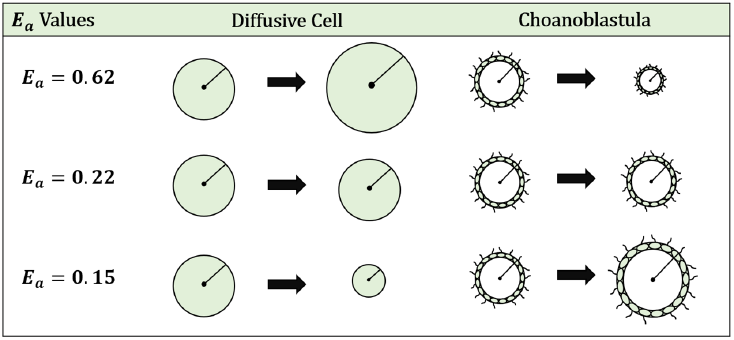
Summary of organism size dependence on temperature for the diffusive cell and the choanoblastula with differing *E*_*a*_ values under constant resource concentration.

The critical radius increases with decreasing nutrient concentration, suggesting organisms using this strategy would have increased in size in response to the environmental changes during Snowball Earth (Fig. 4). Under the best estimates for parameter values, the choanoblastula goes from a minimum radius of approximately 50 *µ*m prior to Snowball Earth to a minimum radius of approximately 10 mm during Snowball Earth. Like the diffusive model, the activation energy *E*_*a*_ impacts the relationship between temperature and organism size. While an activation energy of 0.62 eV results in size decreasing with decreasing temperature, an activation energy below 0.22 eV inverts the relationship (Fig. 7). Regardless of average activation energy, the choanoblastula would have increased in size during Snowball Earth due to the drop in resource availability.

## IV. Discussion

### A. Ecological Changes During Snowball Earth

Changes in temperature, inorganic nutrient concentrations, and light availability had major impacts on the existing organisms and broader ecosystem. The exponential dependence of metabolic rate on temperature caused the primary producer metabolic rates to decrease with temperature, slowing productivity. This decrease is further exacerbated by the physiological and ecological impacts caused by the physical changes accompanying the onset of Snowball Earth glaciations including reduced light under sea ice, higher viscosity, and lower diffusivity. Under the most conservative assumption that primary producer size and population did not change, NPP would still decrease by at least two orders-of-magnitude (Fig. 2a). When the impacts of both nutrient concentration and temperature are considered, that decrease varies between 2.5-4.5 orders of magnitude (Figs. 2b-d).

A reduction in NPP of this magnitude would pose a significant hurdle for heterotrophs, leading to an increase in competition for the remaining resources. This increase in competition was a significant evolutionary driver, which may help to explain why multiple multicellular lineages appeared in this time frame. The diverging response of the two modeled organisms show two possible evolutionary paths. Heterotrophic eukaryotes in the Cryogenian were forced to either get smaller and compete with prokaryotes better suited to the diffusive regime, or become larger, more complex, and multicellular. These observed alternative strategies help explain why some, but not all, eukaryotes evolved multicellularity during this time.

### B. Morphological Differences Lead to Different Adaptive Strategies

A key difference between the two presented morphological models is the scaling between organism size and uptake that originates from two mechanistically different uptake strategies. In the diffusive model, uptake scales with organismal radius due to the physics of diffusion constraining its rate (Eq. 6). By becoming motile and entering the ballistic regime, the choanoblastula uptake rate scales with its cross-sectional area (Eq. 9) and its velocity (Eq. 10), which in-turn scales with organism size. This difference means that an increase in size leads to a large increase in uptake for the choanoblastula compared to the diffusive cell.

Bacterial multicellularity is common and diverse with quorum sensing, metabolic division of labor, large size, and spatial structure [44–48]. In particular stromatolites have a deep geological history, potentially extending back to the first fossil evidence of life [49, 50]. As all bacteria are obligatory diffusion specialists, life within a stromatolite is subject to the same physical processes we model for a solitary diffusive cell [51, 52]. Therefore we can make a first order prediction that the effects of Snowball Earth conditions on stromatolites should match the predictions for solitary diffusive cells. This may provide an additional prediction for the decline in stromatolite abundance and size in the late Neoproterozoic prior to the origin and diversification of grazing and bioturbating bilaterian animals [53, 54].

At the size of eukaryotic cells and simple metazoa, the cost of motility becomes vanishingly small, and provides an enormous benefit for maintaining a larger size by increasing resource uptake (Fig. 5b). However, becoming motile is not enough to offset lower resource availability. The hollow morphology is essential, as it reduces the mass-scaling of metabolic cost of the organism by reducing metabolically active volume while maintaining effective surface area for nutrient uptake. This change in scaling is ubiquitous among complex multicellular organisms, as seen in the infamous two-thirds and three-quarter power laws [27].

Together, these adaptations invert the relationship between nutrient uptake and metabolic rate as a function of organism size. For the diffusive cell, metabolic rate increases faster than uptake, constraining the maximum cell size (Fig. 5a). The opposite is true for the choanoblastula, in which faster uptake means that the energy balance defines a minimum size, allowing it to grow larger until other constraints are reached (Fig. 5b)[39].

### C. Adaptation of Activation Energy

Activation energy (*E*_*a*_) is the amount of energy required to reach a transition state and the source of this energy required to drive reactions is typically heat energy from the surroundings. These results show that organismal size responses to changes in temperature are highly sensitive to activation energy (Figs. 3 and 4). Activation energies vary significantly across life on Earth [55], although much research assumes an average value (0.62 eV, [56]); assuming this value in our models (and thus constraining the relationship between metabolic rate and resource uptake to a specific regime) suggests that diffusive cells must get larger at lower temperatures and the choanoblastula organisms must get smaller (Fig. 7).

However, given the range of measured activation energies, and the fact that unicellular organisms commonly display lower average energies [55], it is necessary to consider differential relationships between metabolic rate and nutrient uptake. The metabolic activation energy emerges from the average activation energies of the underlying enzyme-catalyzed reactions that fuel the organism’s metabolism. Over the 50 million-year glacial period, it is possible that organisms were selected to have lower activation energies in order to maintain their metabolisms at lower temperatures. At an activation energy of 0.22 eV, the body size for both morphological models no longer varies with temperature, and the body size-temperature relationship becomes inverted for both models when the activation energy is less than 0.22 eV. These inversions coincide with the difference in slopes of metabolism under each activation energy relative to the nutrient uptake rate (Fig. 6). Determining the adaptability of metabolic activation energy would be an important step to understanding possible evolutionary trajectories in changing climates.

### D. Pre- and Post-Snowball Dynamics

The paths taken through temperature-resource concentration space during the onset and termination of the Cryogenian glacial periods are important to consider in order to understand the evolutionary trajectories of the existing organisms. Given that primary production decreases due to decreasing temperature and PAR availability, it is likely that temperature decreased faster than resource availability during glacial onset. This trajectory causes diffusive cells to initially grow, reaching their maximum predicted size (*∼* 10^*−*5.2^) while the choanoblastula reach their minimum (*∼* 10^*−*4.5^) (Fig. 8 arrow 1). This places the two modeled organisms in a remarkably similar size range, with radii less than an order of magnitude apart, and at around 10 *µ*m, approximately the size of a modern *Chlamydomonas* [57] or *Salpingoeca* cell [58]. Then, as resource concentrations begin to drop, the organisms’ evolutionary pathways diverge as the diffusive cell is forced to shrink and the choanoblastula grows (Fig. 8 arrow 2).

**Fig. 8.**
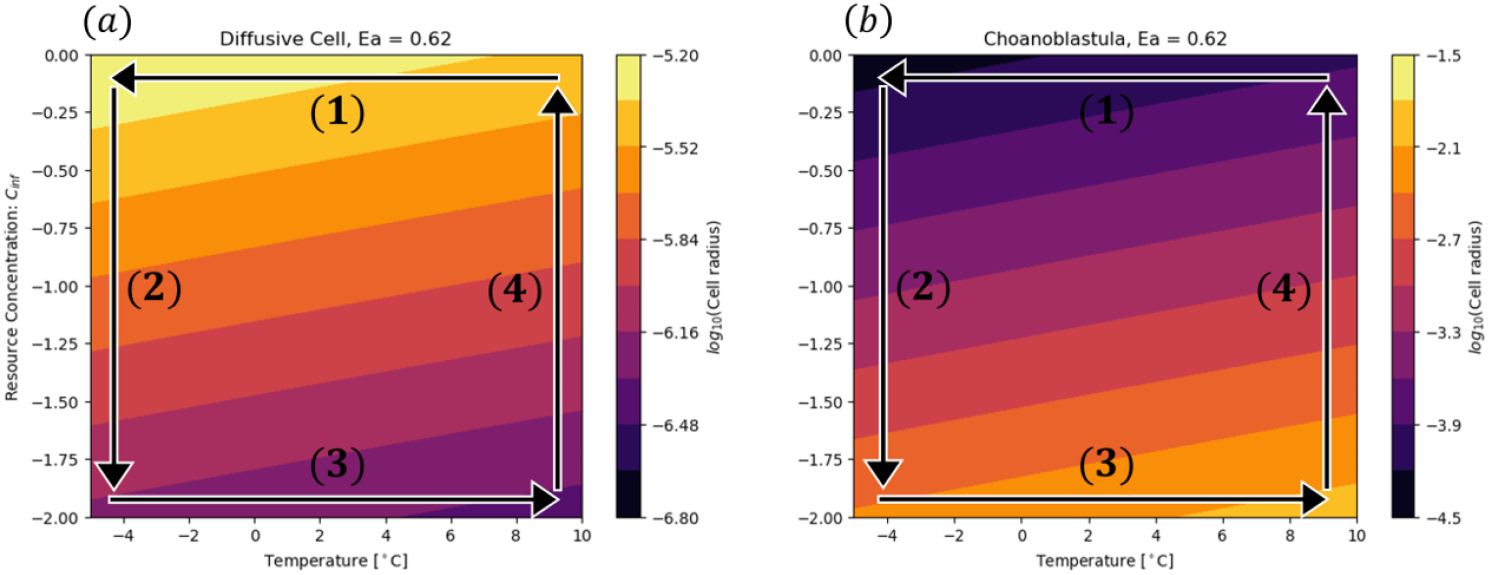
*log*_10_(*radius*) of (*a*) the diffusive cell and (*b*) the choanoblastula, shown as contour plots as functions of temperature and resource concentration. Labeled arrows represent possible trajectories in temperature-resource concentration space for the onset (arrows 1 and 2) and termination (3 and 4) of Snowball Earth.

Following Snowball Earth glaciations, temperature and resource availability increased. Like the onset, it is likely that temperature rebounded before resource concentrations rose. As temperature increased and NPP rates had not yet recovered, choanoblastula would continue to get larger, reaching the maximum predicted size, as the diffusive cell reaches its minimum (Fig. 8 arrow 3). As resource concentrations rise, the model predicts that the choanoblastula would shrink and the diffusive cell would grow (Fig. 8 arrow 4). However this larger size, accompanied by a now increasing amount of resources and faster metabolic rates could allow for new ecological strategies such as predation to develop, allowing the organism to maintain its size as resource availability continues to increase.

These new ecological selective pressures help to explain the rapid proliferation of macroscopic fossils and early metazoan lineages that appear shortly after the end of the glaciations in the Ediacaran.

## V. Conclusions

The only proposed hypothesis for why eukaryotic lineages more readily evolve complex multicellularity is that mitochondria endow eukaryotes with more energetic power which leads to more genes, and consequently more complexity [59]. Given that eukaryotes likely evolved nearly 2 billion years ago [60] and maintained a thriving ecosystem [61], why did it then take over 1 billion years for Eukarya to evolve complex multicellularity? This significant lag between the gain of mitochondria and the evolution of complex multicellularity is not well explained by Lane’s hypothesis. The results above provide an alternative hypothesis that not only explains the timing of the origins of multicellularity but also why bacteria and eukaryotes have such different styles of multicellularity.

Our finding that the metabolic scaling and mechanism of resource acquisition structures the adaptive strategies that emerge during cold, highly viscous, and low nutrient conditions that occur during global glaciations provides a possible mechanism for why bacteria and eukaryotes differ in the nature of their multicellularity. The Cryogenian glaciations therefore provided an opportunity for multicellular eukaryotes to have a selective advantage that bacteria do not share. The need for an environmental trigger helps explain the 1 billion year lag between eukaryogenesis and the appearance of complex multicellular organisms.

The Snowball Earth glaciations may be necessary to provide an opportunity for multicellular eukaryotes to have an adaptive advantage, but they may not be fully sufficient. The eukaryotic-style “always on” gene regulation [62] likely is needed to evolve the more developmentally structured phenotypes needed for multicellularity. Maintaining consistent morphology when reproducing is essential for optimization of size-metabolism scaling and size in response to environmental conditions.

## VI. Acknowledgements

We thank the NSF “RCN for Exploration of Life’s Origins” for facilitating our conversations through a working group hosted at the Santa Fe Institute (NSF Grant 1745355).

