## Appendix for "Physical constraints during Snowball Earth drive the evolution of multicellularity"

### I. Nutrient-Metabolism Model Derivation

To quantify the impact of changing temperature and resource availability on organism morphology, an energy balance model was constructed. The inequality states that the rate of resource uptake ($U$) must be greater than or equal to the rate of energy use by the organism's metabolism ($B$):

$$U \geq B \tag{1}$$

Using this framework, we can then construct mechanistic models of uptake and metabolism in terms of temperature, morphology, and resource concentrations.

#### A. Non-motile Diffusive Model

*1) Resource Uptake Model:* We began by modeling total resource flux into the organism. Here we model a spherical, non-motile cell with a radius $a$ that relies on diffusion for resource uptake (Fig. 1a). Resource uptake is therefore determined by the diffusion equation:

$$\frac{\partial C}{\partial t} = D\frac{1}{r^2}\frac{\partial}{\partial r}\left(r^2\frac{\partial C}{\partial r}\right) \tag{2}$$

Where $C$ is the resource concentration, $D$ is the diffusion constant, $r$ is the radial distance from the center of the cell, and $t$ is time. If we assume that resource concentration is at a steady state, we get the following:

$$\frac{\partial C}{\partial t} = 0 = \frac{1}{r^2}\frac{\partial}{\partial r}\left(r^2\frac{\partial C}{\partial r}\right) \tag{3}$$

Next, to solve for resource concentration $(C)$ as a function of distance from the cell's center $(r)$, we assume that the resource concentration far away from the cell approaches a constant $(C_\infty)$ and the cell takes up all resources at its membrane:

$$\lim_{r\to\infty} C(r) = C_\infty, \qquad C(a) = 0 \tag{4}$$

Using these two conditions, we can solve the partial differential equation for resource concentration as a function of the distance from the cell:

$$x = r^2\frac{\partial C}{\partial r} \tag{5}$$

$$\frac{x}{r^2}\partial r = \partial C \tag{6}$$

$$\int \frac{x}{r^2}dr = \int dC \tag{7}$$

$$B - \frac{x}{r} = C \tag{8}$$

Where $B$ is a constant of integration.

At $r = \infty$:

$$\lim_{r\to\infty} C(r) = B - \frac{x}{r} = B - 0 \Rightarrow B = C_\infty \tag{9}$$

At $r = a$:

$$C(a) = C_\infty - \frac{x}{a} = 0 \Rightarrow x = C_\infty a \tag{10}$$

Resulting in:

$$C(r) = C_\infty \left(1 - \frac{a}{r}\right) \tag{11}$$

Next, to calculate the organism's resource uptake, we use Fick's Law of Diffusion [1]:

$$J(r) = -D\frac{\partial C}{\partial r} = -D\frac{\partial}{\partial r}\left(C_\infty \left(1 - \frac{a}{r}\right)\right) \tag{12}$$

$$J = DC_\infty \frac{a}{r^2} \tag{13}$$

Where $J$ is the flux per unit area at radius $r$. The organism's resource uptake $(U)$ is equal to the resource flux integrated over the surface area of the organism $(A)$ [2]:

$$U = \int JdA = 4\pi DaC_\infty \tag{14}$$

The diffusivity $(D)$ is given by the Stokes-Einstein equation [3]:

$$D = \frac{K_B T}{6\pi\eta R} \tag{15}$$

Where $K_B$ is Boltzmann's constant, $T$ is temperature, $\eta$ is voscosity, and $R$ is the Stokes radius of the resource. We can define viscosity as a function of temperature using the Vogel equation [4]:

$$\eta = \eta_0 e^{\frac{A}{T-C}} \tag{16}$$

Using these relations, we can define the organism's nutrient uptake $U$ as a function of its radius $a$, resource concentration $C$, and temperature $T$:

$$U(a, T, C) = \frac{2}{3}\frac{K_B T}{\eta_0 r} e^{\frac{-A}{T-C}} aC_\infty \tag{17}$$

*2) Metabolic Model:* To model metabolism, we can turn to the Metabolic Theory of Ecology (MTE), which relates an organism's metabolic rate $(B)$ to its temperature $(T)$ and its mass $(M_i)$ [5]:

$$B = b_0 e^{\frac{-E_a}{KT}} M_i^\alpha \tag{18}$$

where $E_a$ is an average activation energy for all metabolic reactions, $b_0$ is a constant, $K$ is Boltzmann's constant, and $\alpha$ is a power-law scaling term. We include a reference temperature $T_0$ to adjust the scaling relationship between metabolism and temperature:

$$B = b_0 e^{\frac{E_a(T-T_0)}{KTT0}} M_i^\alpha \tag{19}$$

To relate the organism's size between its resource uptake, which is dependent on its radius $(a)$, and its metabolic rate, which is dependent on its mass $M_i$, we assume that mass is proportional to volume:

$$M = \rho_c V = \frac{4}{3}\pi\rho_c a^3 \tag{20}$$

Where $V$ is the organism volume and $\rho_c$ is its average density. Incorporating this definition into equation 18, metabolic rate is defined as

$$B = B_0 e^{\frac{E_a(T-T_0)}{KTT0}} a^{3\alpha} \tag{21}$$

where $B_0 = b_0\left(\frac{4}{3}\pi\rho_c\right)^{\alpha}$. Combining the metabolic and resource uptake model into the original inequality gives:

$$\frac{2}{3}\frac{K_B T}{\eta_0 r}e^{\frac{-A}{T-C}}aC_{\infty} \geq B_0 e^{\frac{E_a(T-T_0)}{KTT0}}a^{3\alpha} \tag{22}$$

Solving for organism radius $a$ results in:

$$a \leq \left(\frac{2}{3}\frac{K_B T}{\eta_0 r}e^{\frac{-A}{T-C}}C_{\infty}\frac{e^{\frac{-E_a(T-T_0)}{KTT0}}}{B_0}\right)^{\frac{1}{3\alpha-1}} \tag{23}$$

#### *B. Motile Choanoblastula Model*

As an alternative to the non-motile single cell, we model a hypothetical organism that is motile and hollow (Fig. 1b). Like the non-motile diffusive model, we construct an energy balance between the organism's resource uptake and metabolic rate. Here, we add a work term $(W)$ to the metabolic rate to represent the energetic cost of motility:

$$U \geq B + W \tag{24}$$

*1) Resource Uptake Model:* Rather than relying on diffusion to uptake resources, this organism is motile and encounters resources ballistically [6]. Because the organism is pushing itself through the water and directly intercepting resources, its uptake $(U)$ scales with its cross-sectional surface area $(\pi a^2)$, its velocity $(v)$, and the resource concentration $(C_{\infty})$:

$$U = \pi a^2 v C_{\infty} \tag{25}$$

In order to determine the velocity of the organism, we model it based on power laws that relate velocity $(v)$ to organismal radius $(a)$ and the viscosity of the environment $(\eta)$ [7]:

$$v = \beta a^b \eta^{-m} \tag{26}$$

Where $\beta$ is a proportionality constant, and $b$ and $m$ are scaling coefficients. Estimates of $b$ range from 0.6 to 1, with a value of 0.79 determined by Kiørboe [6]. Similarly, estimates of $m$ range from 0.4 to 4 depending on the species [7]. Expanding equation 26's definition of viscosity using the Vogel equation (equation 16) and using it as a model of velocity in the motile uptake model (equation 25) results in the complete resource uptake model for the motile organism:

$$U = \pi C_{\infty} a^{2+b}\beta\eta_0^{-m}e^{\frac{-mA}{T-C}} \tag{27}$$

*2) Metabolic Model:* The motile organism's metabolic energy expenditure is modeled as the sum of its basal metabolic rate, modeled using the Metabolic Theory of Ecology [5], and a motility cost $(W)$:

$$B = B_0 e^{\frac{E_a(T-T_0)}{KTT0}}M^{\alpha} + W \tag{28}$$

Assuming that the organism exists at a low Reynold's number [7], the cost of motility can be modeled using Stokes' Law for the drag force of a sphere moving through viscous fluid [8]:

$$W = qv^2, \qquad q = 6\pi\frac{a\eta\rho}{\epsilon} \tag{29}$$

Where $q$ is a constant determined by viscosity $(\eta)$, the density of the fluid $(\rho)$, and the energetic-locomotion efficiency $(\epsilon)$, representing the amount of kinetic energy produced by the organism relative to the amount of energy spent on locomotion. Once again, we can write viscosity as a function of temperature using the Vogel equation (equation 16). Density varies with temperature as well, further increasing the temperature dependence.

To determine the mass of the organism $(M)$, we use a hollow-sphere morphology with constant cell size (Fig. 1b). While spherical, the model organism consists of a single layer of cells on the surface of the sphere. The mass of the organism is therefore proportional to the average density of the cells $(\rho_c)$, the average volume of each cell $(\frac{4}{3}\pi R^3)$, and the number of cells $(N)$:

$$M = \frac{4}{3}\pi R^3 \rho_c N \tag{30}$$

The number of cells $(N)$ scales with the spherical organism's surface area:

$$N = \frac{A_{sphere}}{A_{cell}} = \frac{4\pi a^2}{\pi R^2} = 4\frac{a^2}{R^2} \tag{31}$$

Plugging these definitions into MTE (equation 19), results in the following definition for basal metabolic rate:

$$B = B_0 e^{\frac{E_a(T-T_0)}{KTT0}} \left(\frac{16}{3}\pi R \rho_c a^2\right)^{\alpha} \tag{32}$$

Unicellular eukaryotic metabolism scales linearly with mass, so we set $\alpha = 1$ [9].

Combining the models for resource uptake, basal metabolic rate, and mobility cost into the original inequality results in:

$$\begin{aligned} \pi C_\infty a^{2+b} \beta \eta_0^{-m} e^{\frac{-mA}{T-C}} \geq\ & B_0 e^{\frac{E_a(T-T_0)}{KTT0}} 4\pi R a^2 \\ & + \frac{6\pi}{\epsilon} \rho \beta^2 a^{1+2b} \left(\eta_0 e^{\frac{A}{T-C}}\right)^{1-2m} \end{aligned} \tag{33}$$

For $0 < b < 1$, the uptake grows at a greater rate in response to increasing cell radius $a$ than the metabolic cost, which drives an increase in organism size in response to decreasing nutrient availability. This scaling factor relates the organism's size to its motility, and there are a range of estimates, from 0.6 to 1 [7]. Here we will use 0.79 [6].

### II. Global Nutrient Model Derivation

To explore how Snowball Earth impacted evolution, it is important to consider its overall impact on the environment and ecosystem. Here an idealized model of the impact of temperature change and ice cover on global primary productivity is derived in order to explore the effect of planetary glaciation on nutrient availability.

Net primary productivity $(NPP)$ is the measure of the total primary productivity of all organisms in a given volume $(V)$ [10]:

$$NPP = \frac{1}{V}\sum_{i=1}^{n_a} \epsilon P_i \tag{34}$$

Here, $P_i$ represents the productivity of producer $i$, $\epsilon$ is the efficiency of converting light to chemical energy, and $n_a$ is the number of primary producers in the volume $V$. Primary production depends on light availability and the metabolic rate of the producers. The impact of light availability $(I)$ on primary production can be modeled using the Michaelis-Menten equation [10], and metabolism can be modeled using the MTE [5]:

$$NPP = \frac{1}{V}\epsilon p_0 e^{\frac{-E_a}{KT}} \frac{I}{I + K_I} \sum_{i=1}^{n_a} M_i^{\alpha_a} \tag{35}$$

Here, $I$ is the photosynthetically active radiation, measured in moles of photons per square meter per day. $K_I$ is the half-saturating term of the Michaelis-Menten equation, $M_i$ is the mass of producer $i$, and $p_0$ is a normalization constant independent of body size [10].

As ice sheets encroached and covered the majority of the planet's oceans, light availability will drop. Additionally, the impact of temperature on metabolic rate will further slow primary productivity.

To model the impact of both temperature and inorganic nutrient availability, we assume that as Snowball Earth conditions set in, nutrient availability drops by half, modeled by the linear equation $\Delta C_{inf}(T) = \frac{1}{60} * (T - 268.15) + 0.5$.

#### A. *Diffusion Limited Producers*

Primary production depends on the mass $(M_i)$ of each producer. Assuming the producers are non-motile and uptake the nutrients required for photosynthesis through diffusion, the non-motile diffusive model of organisms can be used to model how the primary producers will respond to temperature change. If we assume all producers are the same size, and have the same metabolic scaling factor $\alpha$, we can replace the summation term in equation 35 with $n_a M^{\alpha}$:

$$NPP = \frac{1}{V} \epsilon p_0 e^{\frac{-E_a}{KT}} \frac{I}{I + K_I} n_a M^{\alpha} \tag{36}$$

Next we assume that the producers are spherical and their mass scales linearly with their volume:

$$NPP = \frac{1}{V} \epsilon P_0 e^{\frac{-E_a}{KT}} \frac{I}{I + K_I} n_a a^{3\alpha} \tag{37}$$

Where $P_0 = \frac{4}{3}\pi p_0 \rho_c$. With NPP defined in terms of organism radius, we can use the non-motile diffusive model (equation 23) to model how primary producer's physiology will respond to changes in temperature.

### III. Parameter Values

#### A. *Non-motile Diffusion Model*

| Parameter/Variable | Name | Unit | Value | Source/Reference |
|---|---|---|---|---|
| $a$ | Organism Radius | $[m]$ | Variable | |
| $T$ | Temperature | $[^{\circ}C]$ | Variable, [-5,10] | [7, 11, 12] |
| $C_{\infty}$ | Nutrient Concentration | $[M]$ | Variable | |
| $K_B$ | Boltzmann's Constant | $[J/K]$ | $1.38 * 10^{-23}$ | [13] |
| $r$ | Stokes Radius of Resource | $[m]$ | $3.8 * 10^{-10}$ | [14] |
| $\eta_0$ | Viscosity Constant (of water) | $[mPa * s]$ | 0.02939 | [15] |
| $A$ | Vogel Eqn. Constant 1 (of water) | $[K]$ | 507.88 | [15] |
| $C$ | Vogel Eqn. Constant 2 (of water) | $[K]$ | 149.3 | [15] |
| $C_{yield}$ | Energetic Yield | $[\frac{J}{mol}]$ | $1.4 * 10^{6}$ | [16] |
| $B_0$ | MTE Normalization Constant | $[J/s]$ | $e^{19.21}$ | [5] |
| $E_a$ | Activation Energy | $[eV]$ | 0.62 | [5] |
| $K$ | Boltzmann Constant | $[\frac{eV}{K}]$ | $8.617 * 10^{-5}$ | |
| $T_0$ | Reference Temperature | $[^{\circ}C]$ | 20 | [5, 17] |
| $\alpha$ | Metabolic scaling factor | - | [0.75,1] | [7, 9] |

#### B. *Choanoblastula Model*

| Parameter/Variable | Name | Unit | Value | Source/Reference |
|---|---|---|---|---|
| $v$ | Velocity of organism | $[m/s]$ | $\beta a^b \eta^{-m}$ | [7] |
| $\beta$ | Speed normalizing constant | - | 1 | [7] |
| $b$ | Size-speed scaling factor | - | [0.5,1] | [7, 18] |
| $m$ | Viscosity-speed scaling factor | - | 1 | [7] |
| $R$ | Cell Radius | $[m]$ | Variable | |
| $\epsilon$ | Energetic efficiency of motility | $[J/J]$ | 0.01 | [19] |
| $\rho$ | Fluid density | $[\frac{g}{ml}]$ | 1 | |
| $B_0$ | MTE Normalization Constant | $[J/s]$ | $e^{15}$ | [5] |
| $E_a$ | Activation Energy | $[eV]$ | 0.62 | [5] |
| $\alpha$ | Metabolic scaling factor | - | 1 | [7, 9] |

#### C. Global NPP Model

| Parameter/Variable | Name | Unit | Value | Source/Reference |
|---|---|---|---|---|
| $\epsilon_I$ | Photosynthetic efficiency | $[I/I]$ | 0.1 | [10] |
| $V$ | Volume | $[m^3]$ | - | |
| $I$ | Photosynthetically active radiation | $[\frac{mol*photon}{m^2 s}]$ | Variable | [10] |
| $n_a$ | Number of autotrophic cells | - | $10^5$ | |
| $M_i$ | Mass of autotroph | [kg] | Variable | |
| $\alpha_a$ | Autotroph's metabolic scaling factor | - | 1 | [9] |
| $p_0$ | Production Constant | - | $e^{-11.28}$ | [10] |
| $K_i$ | Production Half Saturation | $[\frac{mol*photon}{m^2 s}]$ | 1.51 | [10] |

### IV. Scripts

All Python Jupyter scripts for the models, parameters, and included figures are open-source and accessible at https://github.com/wwcrockett/SnowballEarthNotebooks.
